# AAV2-mediated intravitreal delivery of exon-specific U1 snRNA rescues optic neuropathy in a mouse model of familial dysautonomia

**DOI:** 10.1101/2025.08.21.671454

**Authors:** Anil Chekuri, Krishnakanth Kondabolu, Emily G. Kirchner, Swanand Koli, Matthew Chagnon, Drenushe Krasniqi-Vanmeter, Max E. Stern, Jessica Bolduc, Giulia Romano, Franco Pagani, Luk H. Vandenberghe, Elisabetta Morini, Susan A. Slaugenhaupt

## Abstract

Familial dysautonomia (FD) is a rare autosomal recessive neurodegenerative disorder caused by a splicing mutation in the *ELP1* gene. It predominantly affects the sensory and autonomic nervous systems, with progressive vision loss due to optic neuropathy being a universal and debilitating symptom. Retinal pathology in FD involves progressive thinning of the retinal nerve fiber layer (RNFL), resulting from the degeneration of retinal ganglion cells (RGCs). Notably, FD-associated vision loss has a postnatal onset, offering a critical window for therapeutic intervention before severe visual impairment develops in adolescence. Currently, no approved treatments exist to prevent or reverse vision loss in FD. In this study, we present a novel RNA-based therapeutic approach targeting *ELP1* pre-mRNA splicing in the retina. We engineered exon-specific U1 small nuclear RNAs (ExSpeU1s) to enhance inclusion of exon 20 in the mutant *ELP1* transcripts in the retina, thereby restoring full-length ELP1 expression. Delivery of ExSpeU1 via adeno-associated virus serotype 2 (AAV2) to the retina improved *ELP1* splicing, rescued RGC loss, and visual function in an FD mouse model. These findings highlight ExSpeU1-mediated splicing correction as a promising therapeutic approach for treating optic neuropathy in FD, offering potential to preserve vision and improve quality of life for patients.

## Introduction

Familial dysautonomia (FD) is a rare congenital sensory and autonomic neuropathy characterized by symptoms such as reduced sensitivity to pain and temperature, early feeding difficulties, cardiovascular instability, and gait ataxia. The average life expectancy for affected individuals is approximately 40 years ^1–5^. FD affects individuals of Ashkenazi Jewish (AJ) ancestry, with a carrier frequency of 1:32 among AJ and 1:17 among AJ of Polish descent^6^. Due to its prevalence of <1/50,000 in the normal population^7^, FD is considered an ultra-rare disease. In addition to their sensory and autonomic deficits, FD patients also suffer from progressive blindness, which onsets in the second decade and severely impacts their quality of life^8–10^. Major ocular defects include decreased visual acuity, loss of central vision, and temporal optic nerve pallor^8–11^. The retinal abnormalities observed in FD patients are similar to other well-characterized optic neuropathies such as Leber hereditary optic neuropathy (LHON) and dominant optic neuropathy (DON)^9,10^. Patients with FD show a significant reduction in the thickness of the retinal nerve fiber layer (RNFL) due to the death of RGCs^8–10^. FD patients suffer from progressive optic neuropathy beginning as early as five years of age^8–10^ with a slowed progression until approximately 15-20 years, followed by a significant visual decline resulting in patients becoming visually impaired by their second decade of life^11^. Since FD patients have diminished proprioception, this visual loss severely affects patient mobility, leading to frequent falls, further compromising their quality of life. The relatively slow rate of progression of visual dysfunction in FD patients provides an excellent therapeutic window for treatment. Therapeutic intervention to treat optic neuropathy at the early stages will significantly enhance the quality of life of FD patients.

FD is caused by a T to C nucleotide change in the 5’ splice site of intron 20 of the elongator acetyltransferase complex subunit 1 (*ELP1,* also known as *IKBKAP*). This mutation leads to skipping of *ELP1* exon 20, resulting in tissue-specific mis-splicing and reduction of ELP1 protein predominantly in the central and peripheral nervous system^1,2,5^. ELP1 plays a major role in catalyzing protein translation, tRNA modification, transcriptional elongation, and cytoskeletal remodeling^12–14^. Mouse models of FD have been instrumental in understanding the retinal pathology characteristic of this neurodegenerative disease^15,16^. Two different conditional knockout mouse models have been previously studied to understand the selective loss of RGCs in the FD retina. In one mouse line, *Elp1* is deleted from neurons in both the central and peripheral nervous system (*Tuba1a-cre; Elp1^flox/flox^*)^16,17^; in the other, *Elp1* is deleted specifically in retinal cells using a *Pax6-cre* (*Pax6-cre;Elp1^flox/flox^)*^15^. In both models, RGCs are lost progressively and postnatally, indicating a cell-autonomous requirement of Elp1 in RGCs for their survival but not development. In addition to these models, we have generated a phenotypic humanized FD mouse model, *TgFD9; Elp1*^Δ20*/flox*^ which carries the human *ELP1* transgene with the FD major mutation and expresses very low amounts of the endogenous *Elp1* gene^18,19^. This mouse reveals progressive loss of the RNFL and ganglion cell layer (GCL), as observed in FD patients, while recapitulating the same tissue-specific skipping of *ELP1* exon 20^18–20^.

Several therapeutic strategies show promise in rescuing RGC loss in mouse models of FD^5,20–25^. Splice modulatory compounds (SMC), which target defective splicing, have been shown to restore RNFL and GCIPL (ganglion cell inner plexiform layer) thickness and preserve RGC counts in FD mice. SMC offers the advantage of systemic delivery, potentially addressing both retinal and systemic defects, and can be administered with relative ease, most likely through oral dosing^20^. However, their off-target effects must be carefully evaluated before clinical translation. Gene replacement therapy for *ELP1* has also been shown to rescue RGC loss in FD, but challenges remain, including potential toxicity from *ELP1* overexpression, which could disrupt protein turnover and cellular homeostasis, and the large size of the *ELP1* gene, which limits precise expression control in target retinal cells. An alternative approach involves antisense oligonucleotides (ASOs) as splice modifiers. ASOs have been shown to significantly correct *ELP1* splicing in transgenic FD mice lacking overt phenotypic defects. However, their efficacy has yet to be tested in a phenotypic model of FD.

U1 small nuclear RNA (U1 snRNA) is a core component of the spliceosome, responsible for recognizing and binding to the 5′ splice site of pre-mRNA via its conserved 9-nucleotide 5′ tail^5^. The 165-nucleotide U1snRNA adopts a stable secondary structure and forms a complex with U1-specific proteins to initiate spliceosomal assembly. Modified versions of U1 snRNA, known as exon-specific U1 snRNAs (ExSpeU1s), are engineered to contain tailored 5′ tails that bind to intronic sequences downstream of canonical 5′ splice sites^22^. These modifications enable ExSpeU1s to promote exon inclusion and effectively rescue aberrant exon skipping in various splicing disorders^22,26–29^, including cystic fibrosis^30^, hemophilia^27,29,31,32^, and spinal muscular atrophy^26,33^. ExSpeU1-mediated rescue has several advantages and promising therapeutic potential. First, by specifically targeting mRNA, gene expression can be efficiently restored while maintaining endogenous transcriptional regulation, limiting the toxic effects of overexpression that are typical of gene supplementation therapies. Second, the compact size of the ExSpeU1 cassette enables splicing correction of genes whose full-length transcripts exceed the packaging capacity of viral vectors, as is the case for *ELP1*. Recent findings indicate that systemic administration of ExSpeU1 was able to correct *ELP1* splicing and rescue survival and gait in the phenotypic FD mice with overall minimal global changes in *Elp1* splicing ^25^. To evaluate the therapeutic efficacy of ExSpeU1 in specifically correcting the *ELP1* splicing defect in the retina and preventing RGC death in FD, we optimized a novel AAV2-mediated intravitreal delivery approach of FD-ExSpeU1 and evaluated the rescue of RGC survival and function. Intravitreal administration of self-complementary AAV2-ExSpeU1 (scAAV2-FD-ExSpeU1) in a phenotypic mouse model of FD rescues RGC survival. More importantly, FD-ExSpeU1 delivery to the retina significantly improved visual function in FD phenotypic mice. The results of this study provide critical proof-of-concept data for the development of a retina-specific splicing modulatory therapy for FD.

## Materials and methods

### Animals

The methodology for the creation of the transgenic mouse line *TgFD9* carrying the human *ELP1* transgene with FD mutation c.2204+6T>C, and the mouse lines carrying *Elp1^flox,^ and Elp1*^Δ20^ alleles were previously published^19,34,35^. The experimental mice used for this study, *F2:TgFD9; Elp1*^Δ20*/flox*^ were generated by crossing the heterozygous mice for *TgFD9* transgene (*TgFD9^+/-^)* and the homozygous mice for the *Elp1 floxed* allele (*Elp1^flox/flox^*) to generate F1 mice (F1:*TgFD9^+/−^; Elp1^flox^*/^flox^). Pups were genotyped to identify those carrying both the *TgFD9* transgene and the *Elp1^flox^* allele (F1: *TgFD9^+/-^; Elp1^flox/flox^*). These F1 animals were then subsequently crossed with the mouse line heterozygous for the *Elp1*^Δ20^ allele (*E1p1*^Δ20*/+*^) to generate the FD mouse (F2:*TgFD9; Elp1*^Δ20*/flox*^). The F2 mice are referred to as FD mice henceforth in the manuscript. Controls are littermates of the FD mice that carry the transgene but are phenotypically normal because they express the endogenous *Elp1 gene (TgFD9; Elp1^+/+^, TgFD9; Elp1 ^flox/+^, or TgFD9; Elp1*^Δ20*/+*^). The expected Mendelian ratio of the FD mouse with this breeding scheme was 1 in 2 (50%). However, - approximately 60% of FD mice survive postnatally due to underlying systemic abnormalities caused by the reduction of *Elp1*^64^. Control and FD mice have a mixed background, including C57BL/6J and C57BL/6N. All the mice enrolled in the study were negative for the spontaneous rd8 mutation, which causes retinal degeneration in the C57BL/6J strain.

### Production of scAAV2-FD-ExSpeU1-eGFP

AAV vector to produce AAV2 expressing FD--ExSpeU1 was generated by cloning ExSpeU1 (750bp) and eGFP (700bp) expression cassettes into ITR (Inverted Terminal Repeat) containing self-complementary AAV2 backbone plasmid. The eGFP cassette is under the control of the cytomegalovirus (CMV) promoter, which is located upstream. Briefly, AAV2 is produced from endo-free plasmid DNA in a triple transfection (ITR-flanked FD-ExSpeU1-eGFP, packaging, and Adeno-helper plasmids) in HEK293T (ATCC,#CRL-3216) cells using PEI (polyethylenimine). After 3 days, the viral vector is isolated from the combined lysate and cell media via sequential high salt, benzonase (for non-AAV protected DNA digestion) treatment, lysate clearing via high-speed centrifugation, Tangential Flow Filtration (for volume reduction and clearing), Iodixanol gradient ultracentrifugation (for purification and separation of empty vs full particles) and buffer exchange and formulation in PBS via molecular weight cutoff filtration. The preparation underwent titration via digital droplet PCR, and quality control for purity was assessed via SDS-PAGE.

### Intravitreal injections

Experimental and control mice were anaesthetized with an intraperitoneal injection of a ketamine/xylazine mixture (70mg/kg ketamine and 20mg/kg xylazine). Pupils of the mice were dilated using 2.5% phenylephrine and 1% tropicamide. Approximately 0.5% proparacaine is used as a topical anesthetic during the procedure for 1-2 minutes. GenTeal gel was used to keep the eyes moist and avoid corneal dryness and opacities. The body temperature of the mouse was kept stable at 37 ^°^C throughout the procedure. Under the control of a stereo microscope (Discovery.V20, Zeiss), anincision was made into the sclera posterior of the superior limbus using a sterile, sharp 30-gauge (G) needle without touching the lens. A microliter Hamilton syringe attached to a 34G blunt needle was carefully inserted into the same incision, and 1 μl of *scAAV2-FD-*ExSpeU1-eGFP *or* PBS1X (sterile phosphate-buffered saline) was injected slowly into the vitreous using manual pressure. The needle was kept inside the vitreous area for at least 1 minute after the injection to avoid reflux of viral suspension. The right eye is injected with scAAV2-FD-ExSpeU1-eGFP, and the contralateral eye is injected with 1XPBS. After the completion of intravitreal AAV injections, triple antibiotic ointment (neomycin, bacitracin, and polymyxin B) was applied, and mice were injected subcutaneously with the analgesic buprenorphine (0.1 mg/kg).

### Spectral-domain optical coherence tomography (SD-OCT)

To evaluate the retinal structure, we performed *in vivo* imaging of the retina using SD-OCT. The mice were anesthetized by a ketamine/xylazine mixture as described above. Pupils of the mice were dilated using 2.5% phenylephrine and 1% tropicamide. Approximately0.5% of proparacaine is used as a topical anesthetic during the procedure. SD-OCT imaging was performed using a Leica EnvisuR2210 OCT machine. RNFL and GCIPL measurements were performed by 10-layer automated segmentation using the software DIVER (Leica). PBS-injected and scAAV2-FD-ExSpeU1-eGFP-injected eyes of control and FD mice at 6 months of age were analyzed. Each OCT image comprises 100 B-scans, with each B-scan consisting of 1000 A-scans.

### Retinal whole mounts

RGC staining and counting of retinal whole mounts were performed according to the method previously described by Ueki et al^15,16^. Briefly, fixation of the eyes was performed at room temperature for 1 h in 4% PFA (paraformaldehyde), and the eyes were marked with a yellow tissue marking dye on the temporal surface. After fixation, retinas were isolated, with each temporal retina marked with a small cut. Relaxing cuts in the spherical retina were made on all four corners to flatten the retina. Non-specific binding was blocked by incubation with an animal-free blocker containing 0.5% Triton X-100 overnight at 4°C. RGC-specific anti-RNA-binding protein with multiple splicing (RBPMS) antibody (PhosphoSolutions #1830-RBPMS) was applied for 48 hrs. at 4°C. Retinas were incubated with secondary antibody (Thermo Fischer Scientific Alexa Fluor 568 **#** A-11011) for 1 h at room temperature and mounted on slides. Images were acquired with a Leica DMi8 epifluorescent microscope and MetaMorph 4.2 acquisition software (Molecular Devices, San Jose, CA). Whole scans of complete flat-mount samples were obtained at 20X magnification via scan stage and autofocus. The total retinal area across the entire scan was approximately 14 mm^2^. With ImageJ software, the number of RBPMS +ve cells was measured at 1 mm^2^ at 1 mm from the optic nerve head (ONH) at the superior, inferior, temporal, and nasal hemispheres^15^. If a specific square area was damaged as a result of rips or folds in the retina, we counted the RGCs in an adjacent undamaged area.A customized analysis pipeline was developed in CellProfiler to objectively identify and count RBPMS+ve cells based on their characteristic size, intensity, and morphology. The pipeline included steps for image preprocessing, primary object (cell) identification, and intensity-based filtering to distinguish positive signals from background. Cell counts were performed within a defined 1 mm² region of interest positioned 1 mm from the optic nerve head in each retinal quadrant (superior, inferior, nasal, and temporal). The investigator conducting the analysis was blinded to the genotype.

### Histology

Eyes were enucleated and fixed in 4% PFA overnight at 4°C. After a single 1XPBS wash, eyes were cryoprotected in 30% sucrose overnight at 4°C and embedded in optimal cutting temperature compound (Sakura Finetek, Torrance, CA, USA) and cryo-sectioned into 15-μm thin sections. Sections were fixed in dry ice-cold acetone for 15 min and washed with 1XPBS three times. Antigen retrieval was then performed by incubating the sections using the buffer containing 0.1M Tris.HCL pH 8, 50 mM EDTA pH 8.0, and 20 μg/ml proteinase K for 10 min at room temperature. After washing two times in 1XPBS, retinal sections were blocked with an animal-free blocker (Vector Laboratories, Burlingame, CA, USA) containing 0.5% Triton X-100 for 1 h at room temperature. Primary antibody anti-GFP (Green Fluorescent Protein) (Thermo Fisher Scientific, #A10262, CA, USA) was applied and incubated at 4°C overnight in a moist chamber. Sections were washed three times with 1XPBS with 0.1% Triton X-100 and incubated with secondary antibodies (Invitrogen; Jackson Immuno Research, West Grove, PA, USA) for 1 h at room temperature. Sections were mounted with hard-set mounting media containing DAPI (Vector Labs), and fluorescent microscopy was performed. One eye from each mouse was included in this analysis.

### Real-time-qPCR (RT-qPCR)

RT-qPCR detection of the expression of FD-ExSpeU1 was performed using RT-qPCR LightCycler 480 II (Roche). RNA was extracted from the retinas of either PBS-injected or scAAV2-FD-ExSpeU1-eGFP-injected mice using an RNA isolation kit (Qiagen) and treated with DNase. RNA yields were quantified using a NanoDrop ND-1000 spectrophotometer. RT was performed for cDNA synthesis with 500 ng of starting RNA in an 11.5 μL RT reaction. qPCR was performed with iQ-SYBR Green Supermix (Bio-Rad Laboratories) using FD-ExSpeU1-specific primers (forward,5’-ATAGCAAACAGTACAATGC-3’and reverse,5’-CACTACCACAAATTATGCA-3’). The expression levels of FD-ExSpeU1 were directly normalized with the housekeeping gene (*Gapdh*). Quantitative real-time PCR was carried out at the following temperatures for the indicated times: 95 °C (3min);45 cycles of 95 ^°^C (15sec) followed by 60 ^°^C (30 sec); and 98 ^°^C (0.11sec). Data was analyzed with the SDS software.

### RT-PCR analysis of *ELP1* transcripts

Mice were euthanized, and eyes were enucleated in ice-cold 1xPBS. Retinas were dissected on ice under a microscope (Zeiss) and homogenized in ice-cold QIAzol Lysis buffer (Qiagen) with Qiagen tissue Lyser. Similarly, RNA was isolated with QIAzol Lysis Reagent from human fibroblasts treated with scAAV2-FD-ExSpeU1-eGFP, and total RNA was extracted using the standard chloroform extraction procedure. The concentration of the total RNA for each sample was determined with a Nanodrop ND-1000 spectrophotometer. cDNA was synthesized by reverse transcription with ∼0.5 μg of total RNA, Random Primers (Promega), and Superscript III reverse transcriptase (Invitrogen) according to the manufacturer’s instructions. To perform RT-PCR and detect *ELP1* exon 20 inclusion, 100 ng of starting RNA equivalent cDNA was subjected to a PCR reaction with the GoTaq green master mix (Promega,#M7122) and 30 amplification cycles (94°C for 30 s, 58°C for 30 s, and 72°C for 30 s). The human-specific forward and reverse primers spanning exon 19–21 of the *ELP1* 5′-CCTGAGCAGCAATCATGTG−3′ and 5′-TACATGGTCTTCGTGACATC-3′, were used for amplification of both wild-type and mutant isoforms of the human *ELP1 transcript*. The PCR products were resolved on 1.5% agarose gels containing ethidium bromide for detection. The relative amounts of normal and mutant *ELP1* isoforms were quantified using ImageJ software, and the integrated density value for each band was determined and expressed as a percentage as described previously^19^.

### Cell culture and Minigene assay

HEK293T (ATCC, #CRL-3216) cells were cultured in Dulbecco’s modified Eagle’s medium (11995-065, D-MEM, Gibco) supplemented with 10% fetal bovine serum (FBS, 12306C, Sigma) and 1% penicillin/streptomycin (30-009-CI, Corning). HEK293T cells were transfected with FD-minigene and scAAV2-FD-ExSpeU1-eGFP constructs and seeded in 6 wells. The following day, the media was changed to regular growth media. The total RNA was extracted using the standard chloroform extraction procedure 48 hours after transfection to detect the *ELP1* transcripts. To specifically amplify the spliced *ELP1* transcript expressed by the FD minigene, T7 (5′-TAATACGACTCACTATAGGG) and BGHR (5′-TAGAAGGCACAGTCGAGG) primers were used. The PCR products were separated on a 1.5% agarose gel containing ethidium bromide and visualized with the Alpha 2000 Imager Analyzer (Bio-Rad). Band intensities were quantified using ImageJ software. The expected sizes of the amplified products were approximately 480 base pairs for the mutant *ELP1* transcript and 550 base pairs for the wild-type *ELP1* transcript. Patient human fibroblasts carrying the c.2204 + 6T > C mutation in *ELP1* were obtained from *C*oriell Cell Repository (GM04663). Approximately 1 × 10^4^ human FD fibroblast cells were plated in 6-well plates and incubated at 37°C for 12 hr. Cells were washed once with 1XPBS (room temperature) and then infected at 37°C either with phosphate-buffered saline (control) or with scAAV2-FD-ExSpeU1-eGFP vectors at a multiplicity of infection (MOI) of approximately 1,000 vgs/cell. 4 hours after infection, the virus was removed by washing the cells with DMEM/F12 media once, and the cells were supplemented with complete media. After 48 hours of infection, cells were homogenized in ice-cold QIAzol Lysis buffer (Qiagen, #79306), and the total RNA was processed according to the manufacturer’s instructions. The isolated RNA was then subjected to RT-PCR as described above.

### Pattern Electroretinography (PERG)

Mice were anesthetized and mounted on the Celeris PERG system (Diagnosys) platform and kept under dim red light throughout the procedure. A corneal electrode was placed on the right eye, and body temperature was stabilized at 37°C. The visual stimuli were presented at 98% contrast and constant mean luminance of 50 cd/m^2^, spatial frequency: 0.05 cyc/deg, and temporal frequency: 1Hz. A total of 300 complete contrast reversals of PERG were repeated twice in each eye, and the 600 cycles were segmented, averaged, and recorded. The averaged PERGs were analyzed to evaluate the peak-to-trough P1 to N2 amplitude as described previously^36^.

### Optomotor response (OMR)

To assess the visual function in the scAAV2-FD-ExSpeU1-eGFP-treated eye compared to the sham-treated eye, visual acuity in mice was assessed using an automated optomotor response (OMR) machine (OptoDrum, STRIATECH GmbH). The optomotor response is evaluated via a spatial frequency threshold test. The equipment is composed of four computer monitors arranged in a quadrangle, which simultaneously display black and white striped patterns. The mouse is positioned on a pedestal at the center, directly below the camera. Tracking behavior was automatically detected and analyzed by OptoDrum software. This test includes an observer to make sure that all the mice’s behaviour is tracked throughout the experiment. The tracking is recorded only when the mice are sitting still, for a maximum of 20 seconds of still-sitting time. Tracks in the counterclockwise direction are subtracted from the total tracking period, and if the response surpasses the user’s predetermined threshold, the trial is considered positive. For visual acuity, the contrast of 99.27% and rotation speed (12° s −1) were kept constant, and the cycle per degree was adjusted in a preprogrammed staircase method. For the outcome, two positive and three negative trials at the next higher cycle per degree are required. The resulting value representing visual acuity corresponds to the highest spatial frequency observed during the experiment.

### Statistical analysis

Data are presented as mean ± SEM. Statistical significance was determined using a two-tailed unpaired Student’s t-test with False Discovery Rate (FDR) correction for multiple comparisons, or by ordinary one-way ANOVA where appropriate, as indicated in the figure legends. Adjusted value of *p < 0.05 ^∗∗∗^*p < 0.01,*^∗∗∗^*p < 0.001 and*^∗∗∗∗^*p < 0.0001,* was considered statistically significant. All analyses were performed using GraphPad Prism software.

## Results

### Design and development of scAAV2-FD-ExSpeU1-eGFP for retinal delivery

Mutations in the 5′ splice site (5’ss) can compromise U1 snRNA binding, impair spliceosome assembly, and consequently lead to exon skipping, as observed in FD. ExSpeU1s are ∼86-nucleotide modified U1 snRNAs that are designed to bind intronic sequences via a modified 5′ tail. Encoded by a compact ∼700 bp region, they are well-suited for in vivo delivery using AAV. By targeting the 5′ splice site downstream of specific exons, ExSpeU1s can correct aberrant splicing in various cellular and mouse models ^26–30,33,37^. Additionally, they can promote the inclusion of exons that are normally alternatively spliced ^38^. To restore defective splicing, ExSpeU1s have been created with sequence changes that permit targeted binding to intronic sequences downstream of the mutant 5’ ss^26,30,33,39^. Our previous work has explored the use of the minigene system to screen ExSpeU1s that can efficiently correct the *ELP1* exon 20 splicing of C.2204+6 T>C mutation. Using this assay, we have identified Ik10 as the most effective ExSpeU1 snRNA species that can correct exon 20 splicing both in vitro and in vivo^22^. Additionally, we have also shown that AAV9-mediated delivery of ExSpeU1 via intraperitoneal (IP) and intracerebroventricular (ICV) routes leads to significant improvement of gait ataxia, renal and cardiac function in FD mice^25^. Given the recent success in retinal gene therapy and the impact of vision loss in FD patients, we asked whether local delivery of FD-specific ExSpeU1 can correct *ELP1* splicing in the retina and prevent RGC loss in the FD phenotypic mouse *TgFD9; Elp1*^Δ20*/flox*^. To develop a novel ExSpeU1-based approach capable of efficiently and specifically correcting *ELP1* splicing defects in the retina, we have designed a recombinant self-complementary adeno-associated viral (AAV) plasmid vector, (scAAV-FD-ExSpeU1-eGFP), consisting of a sequence encoding ExSpeU1 (U1snRNA) and an enhanced green fluorescent protein (eGFP) reporter gene under the control CMV promoter. These two genes are flanked by two inverted terminal repeat (ITR) sequences to generate a plasmid vector compatible with AAV production. (*Figure 1A*). We chose to express eGFP along with FD-ExSpeU1 to study the retina-specific expression of FD-ExSpeU1. The efficacy of FD-ExSpeU1 in promoting inclusion of *ELP1* exon 20 was first evaluated using our well-established FD minigene splicing assay.^40,41^. A minigene containing *ELP1* exons 19, 20, and 21 and introns 19 and 20, along with the FD splice site mutation, was transfected into HEK293T cells with the plasmid sc*AAV-FD-ExSpeU1-eGFP*. Transfection of increasing concentrations of scAAV-FD-ExSpeU1-eGFP corrected *ELP1* splicing in a dose-dependent manner *(Figure 1B and C, Figure S1*). To deliver FD-ExSpeU1 to the retina and, more specifically, to the RGCs, the retinal neurons that are affected in FD, we have produced a self-complementary AAV2 expressing ExSpeU1 (scAAV2-FD-ExSpeU1-eGFP). AAV2 capsid was used to deliver ExSpeU1 based on its reported tropism for mouse RGCs and its use and tolerance in the human eye^42,43^. To initially test the efficacy of scAAV2-FD-ExSpeU1-eGFP in correcting *ELP1* splicing in vitro, FD fibroblasts obtained from patients (containing c.2204+6T>C mutation in the *ELP1* gene) were transduced with scAAV2-FD-ExSpeU1-eGFP for 48 hours. RT-PCR analysis to detect the *ELP1* transcripts indicated the complete correction of the splicing defect in FD fibroblasts compared to the control untransduced fibroblasts (*Figure 1D, E, and Figure S2*).

**Figure 1:**
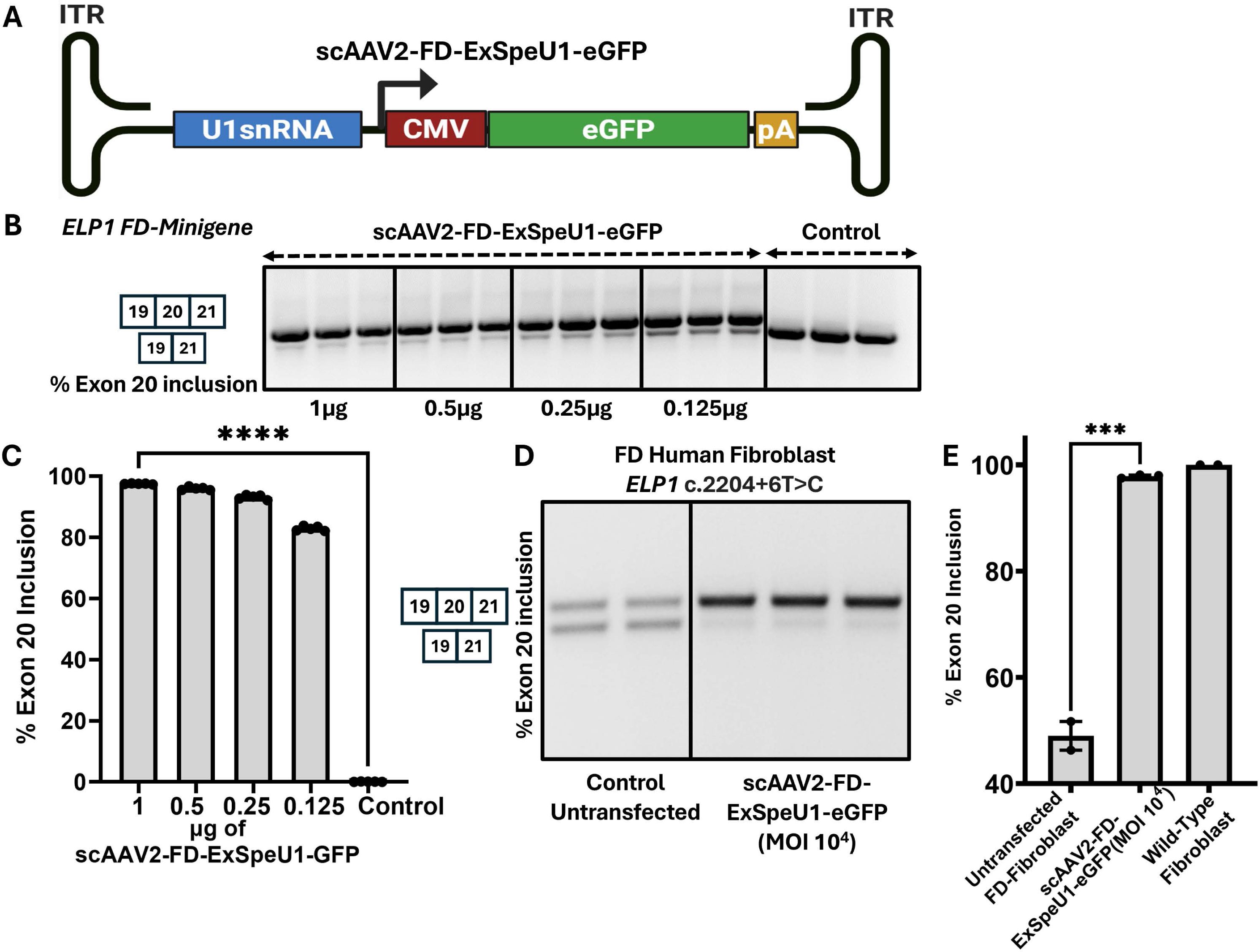
Design and *in vitro* testing of AAV2 vector expressing FD-ExSpeU1. **A.** Schematic of the AAV vector expressing FD-ExSpeU1 and eGFP. FD-ExSpeU1 gene sequence, along with eGFP under the control of cytomegalovirus (CMV) promoter and bovine growth hormone (BGH polyadenylation A) (pA) sequences, was cloned between two inverted terminal sequences (ITRs). **B.** Representative gel picture showing splicing correction of FD minigene in HEK293T cells with increasing concentration of scAAV2-FD-ExSpeU1-eGFP vector *(n=5), (See also Figure S1)*. **C.** Bar graph indicating the quantification of exon 20 splicing correction in FD minigene assay (\*\*\*\**p<0.0001*). **D.** Representative gel picture showing complete correction of *ELP1* splicing in FD patient fibroblasts (*ELP1*c.2204+6T>C) infected with scAAV2-FD-ExpeU1-eGFP, (*See also Figure S2*)*. E.* Bar graph indicating the quantification of exon 20 splicing correction in FD Fibroblast (*ELP1*c.2204+6T>C) transfected with scAAV2-FD-ExpeU1-eGFP. Statistical analysis was performed using ordinary ANOVA to compare the differences between the groups. Adjusted p-values are shown: ^∗∗∗∗^*p < 0.0001,* ^∗∗∗^*p < 0.001*.

### Intravitreal delivery of scAAV2-FD-ExSpeU1-eGFP corrects *ELP1* splicing in the retina

We further aimed to establish the most effective ExSpeU1 delivery system to correct the *ELP1* splicing defect in the retina. We initially employed the asymptomatic *TgFD9* transgenic mice harboring the human *ELP1* transgene with the c.2204+6T>C FD mutation. These mice are phenotypically normal as they express normal levels of endogenous mouse *Elp1*. This is the best model to assess the effect of our ExSpeU1-based approach on *ELP1* splicing. We chose to utilize *TgFD9* mice for this analysis, as phenotypic FD mice are more difficult to generate due to their low survival rate^19^. We administered intravitreal injections of scAAV2-FD-ExSpeU1-eGFP into *TgFD9* mice at postnatal day 21 (P21), delivering 1 × 10¹ viral genomes (VG) in a volume of 1 µL per eye. This dose was selected based on prior studies demonstrating its efficacy and safety as an optimal range for intravitreal delivery in murine models^44,45^. Notably, similar dosing strategies have been employed in clinical trials targeting Leber Hereditary Optic Neuropathy (LHON), further supporting its translational relevance ^45^. We then evaluated the transduction efficacy of scAAV2-FD-ExSpeU1-eGFP 30 days post-injection by staining retinal flat mounts with a GFP-specific antibody. Intravitreal administration of scAAV2-FD-ExSpeU1-eGFP resulted in the expression of eGFP throughout the retina, indicating significant transduction efficiency (*Figure 2A*). The expression of eGFP was predominantly observed in RGCs, as evidenced by the GFP staining of the retinal cross sections obtained from the eyes injected with scAAV2-FD-ExSpeU1-eGFP (*Figure 2B*). In addition, to specifically evaluate the expression of ExSpeU1 in the retinas injected with scAAV2-FD-ExSpeU1-eGFP in FD phenotypic mice, qPCR analysis was performed. A significant expression of ExSpeU1 was detected in injected retinas of FD mice compared to 1XPBS-injected control retinas (*Figure 2C)* (*p<0.053*). We further analyzed the correction of *ELP1* splicing in the retinas injected with scAAV2-FD-ExSpeU1-eGFP, which showed significant correction of the splicing compared to control (*p<0.001*) (*Figures 2D, E and Figure S3*). These results indicate efficient retinal targeting of the scAAV2-FD-ExSpeU1-eGFP vector and significant correction of *ELP1* splicing with intravitreal delivery of ExSpeU1.

**Figure 2:**
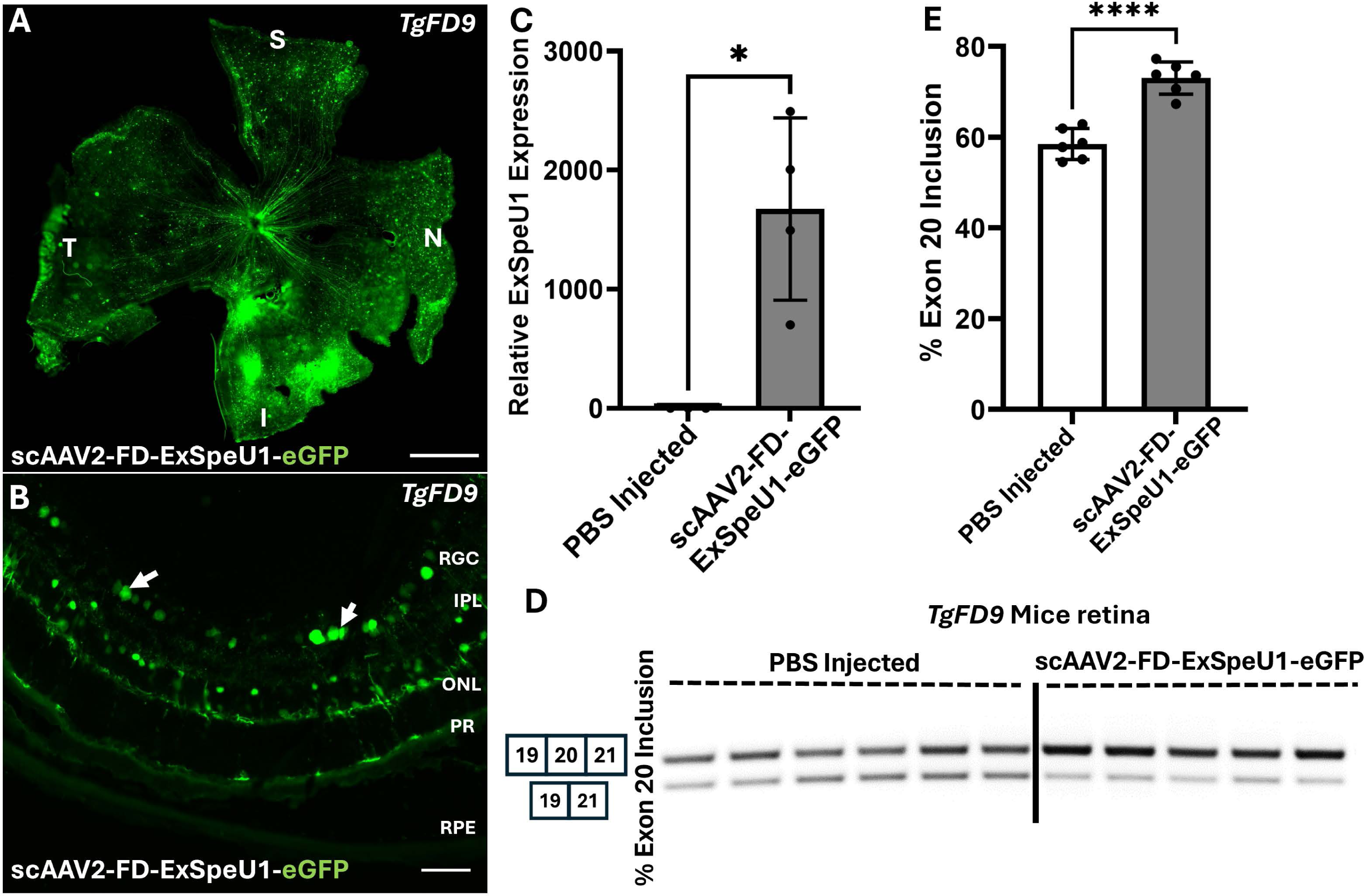
scAAV2-FD-ExSpeU1-eGFP corrects *ELP1* splicing defect in the retina. **A.** Representative image showing retinal flat mount from *TgFD9* mice intravitreally injected with scAAV2-FD-ExSpeU1-eGFP. Staining was performed using a anti-GFP antibody. **B.** Image of retinal cross-section obtained from *TgFD9*mice stained with anti-GFP antibody. The arrow indicates GFP+ve cells in the RGC layer of the retina. **C**. Bar graph representing the relative expression of FD-ExSpeU1 in the retina of FD phenotypic mice injected with scAAV2-FD-ExSpeU1-eGFP compared to PBS-injected control 6 months after injection (*n=4; p<*0.053*). **D**. Representative splicing analysis of human *ELP1* transcripts (both the PCR products from the gel were confirmed by Sanger sequencing), (*See also Figure S3*). **E.** Percent of exon 20 inclusion from scAAV2-FD-ExSpeU1-eGFP (n=6, light grey). ***P< 0.001, two-tailed unpaired Student’s t-test with FDR (False Discovery Rate) correction.

### scAAV2-FD-ExSpeU1-eGFP treatment rescues RNFL loss in FD mice

We have previously shown that FD mice exhibit significant RNFL and ganglion cell inner plexiform layer (GCIPL) loss starting at 3 months, with further reduction at 6 months^18^. The progressive thinning of these retinal layers perfectly recapitulates the disease phenotype observed in FD patients^18^. To test whether delivery of scAAV2-FD-ExSpeU1-eGFP can rescue RNFL and GCIPL thinning, we performed intravitreal injections of scAAV2-FD-ExSpeU1-eGFP at a total dose of 1×10^10^ viral genomes at P30 in the phenotypic FD mice. One eye in FD mice is injected with scAAV2-FD-ExSpeU1-eGFP, and the contralateral eye is injected with 1XPBS (Sham). To evaluate the rescue of RNFL and GCIPL thickness in vivo, we performed a spectral domain-OCT (SD-OCT) analysis. We measured RNFL thickness in the superior, inferior, nasal, and temporal hemispheres of the retinas at 6 months of age in injected mice (*Figure 3A)*. We observed a significant reduction of the RNFL in each region of the PBS-injected retinas (sham) compared to control littermates (central temporal; *p<0.0017,* central nasal; *p<0.0021,* peripheral temporal; *p<*0.0001, and peripheral nasal; *p<*0.0001).

**Figure 3:**
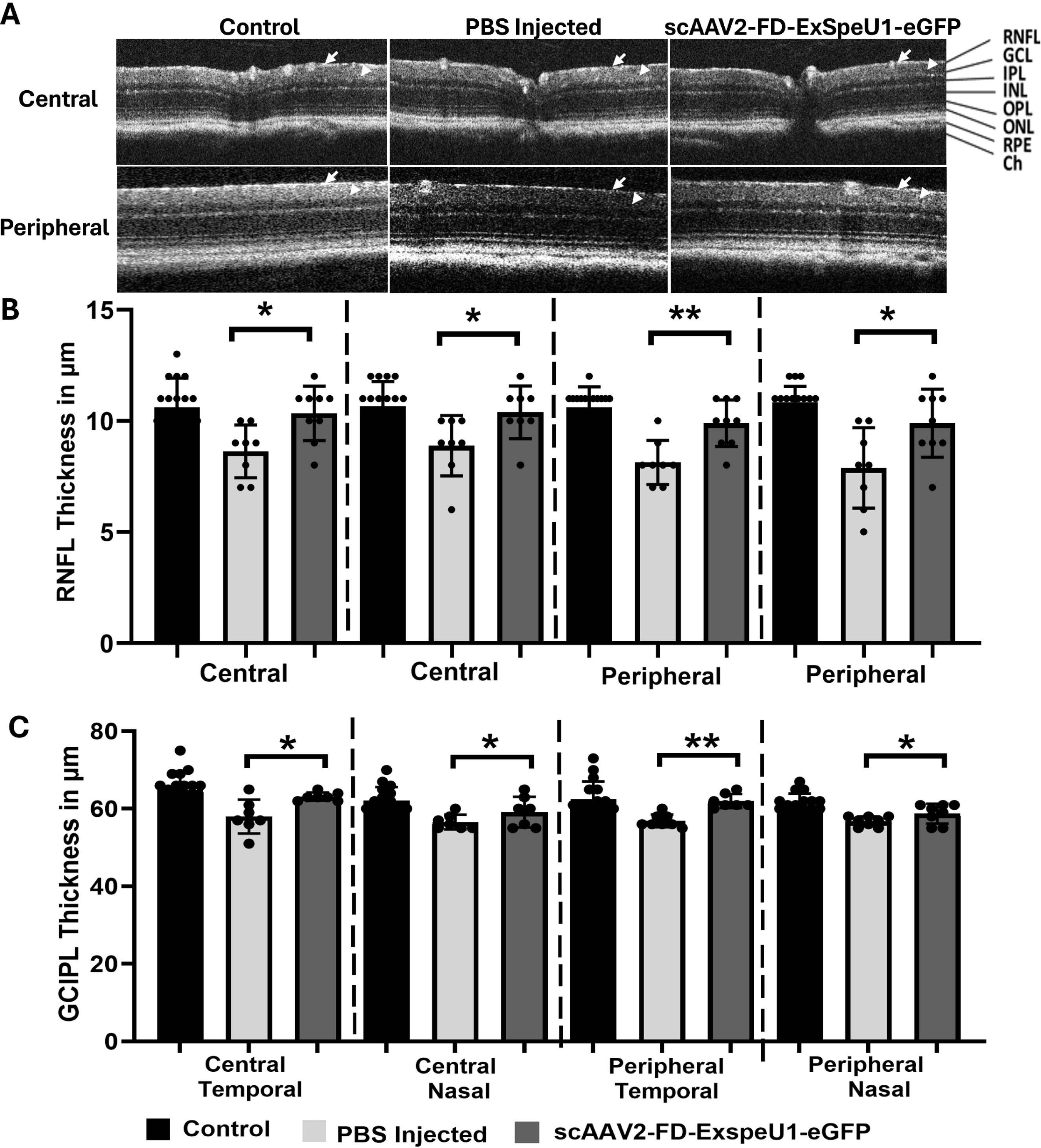
Intravitreal delivery of *scAAV2-FD-ExSpeU1* improves RNFL and GCIPL thickness in FD mice: **A.** Representative image of SD-OCT analysis in FD mice intravitreally injected with scAAV2-FD-ExSpeU1-eGFP six months post-injection. One eye of the FD mice is injected with AAV, and the contralateral eye is injected with 1XPBS. **B.** Measurement of RNFL thickness in both temporal and nasal hemispheres in central and peripheral regions of the retina in control (n=8) and scAAV2-FD-ExSpeU1-eGFP-injected eyes and PBS-injected in FD mice (n=8). **C.** Bar graph representing improvement in GCIPL thickness in scAAV2-FD-ExSpeU1-eGFP-injected FD eyes compared to PBS-injected eyes (n=8). The adjusted p-values are displayed. ^∗^*p < 0.05,* ^∗∗^*p < 0.01* , two-tailed unpaired Student’s *t* test with FDR correction. Data are shown as average ± SEM; each data point represents an individual retina.

Interestingly, retinas injected with scAAV2-FD-ExSpeU1-eGFP showed significant improvement in the RNFL thickness compared to sham (central temporal; *p<0.0107,* central nasal; *p<0.0337,* peripheral temporal; *p<*0.0030, and peripheral nasal; *p<*0.0253) (*Figure 3B*). Similarly, GCIPL thickness was significantly reduced in FD mice compared to control reconfirming our previous findings in FD mice^18^ (central temporal; *p<0.0038,* central nasal; *p<0.0010,* peripheral temporal; *p<*0.028, and peripheral nasal; *p<*0.0010). In addition, GCIPL thickness was improved in retinas injected with scAAV2-FD-ExSpeU1-eGFP in all retinal regions compared to saline-injected retinas (central temporal; *p<0.0104,* central nasal; *p<0.0203,* peripheral temporal; *p<*0.001, and peripheral nasal; *p<*0.0028) (*Figure 3C*). These results indicate that delivery of scAAV2-FD-ExSpeU1-eGFP was able to improve RNFL and GCL thickness, supporting the efficacy of our approach in rescuing retinal nerve fiber in FD mice.

### Delivery of scAAV2-FD-ExSpeU1-eGFP to the retina ameliorates RGC degeneration in FD mice

Progressive death of RGCs postnatally has been shown in well-characterized mouse models of FD^15,16^. Our previous study to characterize retinal phenotype in the FD mouse model indicated a significant loss of RGC at 6 months^18,20^. To further substantiate the findings from the OCT analysis and correlate them with histological rescue, we performed RGC counting in the retinas collected from scAAV2-FD-ExSpeU1-eGFP and PBS-injected FD mice in the superior, inferior, nasal, and temporal regions of the retina by staining the retinal flat mounts with RGC specific marker RNA-binding protein with multiple splicing (RBPMS) (*Figure 4A*). Consistent with our previous observations, the number of RBPMS +ve cells is significantly lower in PBS-treated retinas in all the retinal regions compared to retinas from control littermates (superior; *p<0.0003,* inferior; *p<0.0329,* temporal; *p<0.0471,* and nasal; *p<0.0172*). Importantly, significant rescue of RGC loss was observed in the retinas treated with scAAV2-FD-ExSpeU1-eGFP in the temporal, nasal, superior, and inferior regions of the retina compared to the saline-injected retinas (superior; *p<0.0449,* inferior; *p<0.0473,* temporal; *p<0.0284* and nasal; *p<0.0026*) (*Figure 4B*). Together, these results demonstrate that intravitreal delivery of ExSpeU1 can halt the progression of RGC loss observed in FD mice.

**Figure 4.**
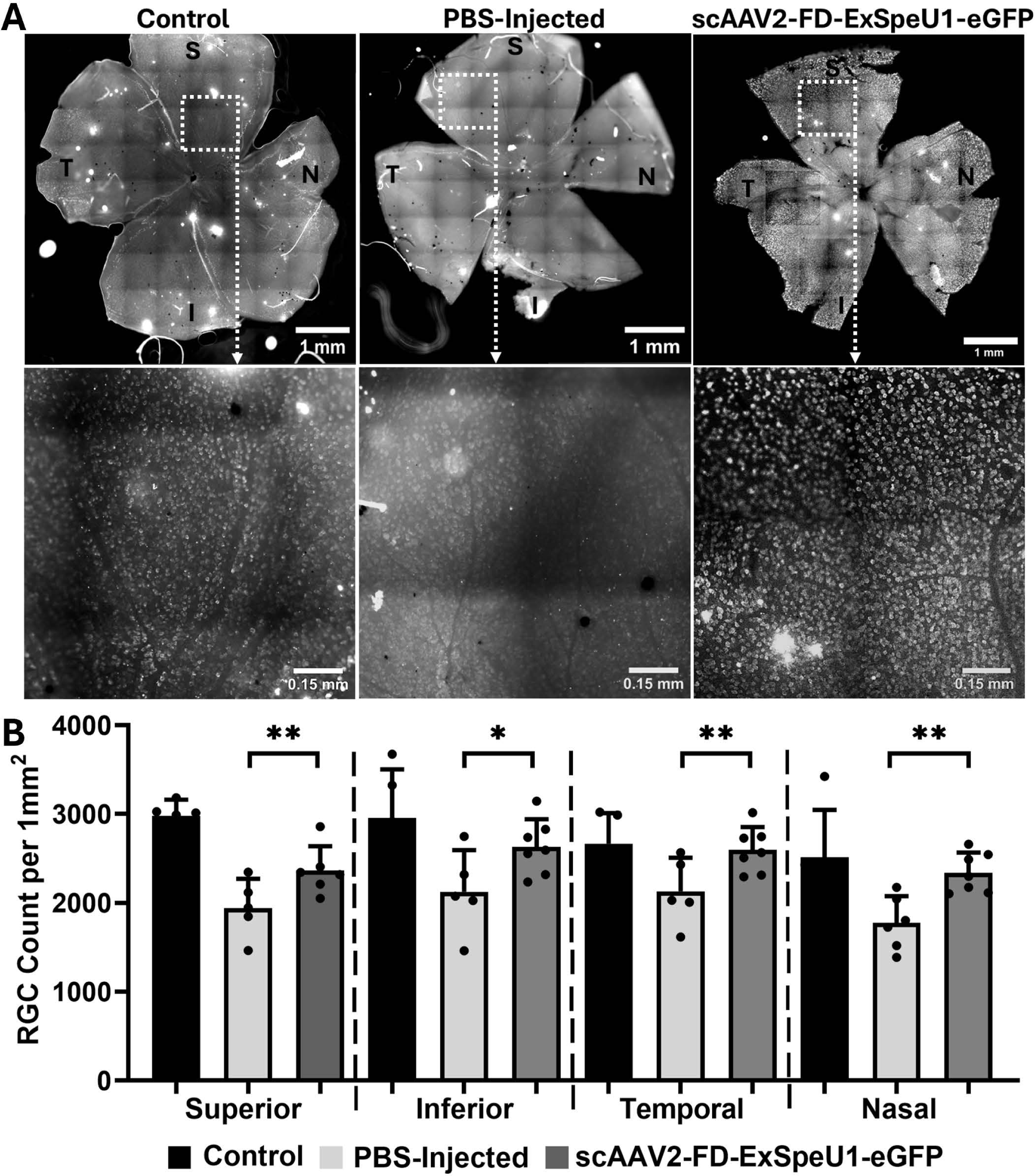
scAAV2-FD-ExSpeU1-eGFP ameliorates RGC degeneration in FD mice. A. Representative flat-mount images of retinas from control mice, PBS-injected FD mice (Sham), and scAAV2-FD-ExSpeU1-eGFP–injected FD mice at 6 months of age. Scale bar: 1 mm. RBPMS+ve RGCs were quantified in the superior, inferior, nasal, and temporal quadrants at 1 mm from the optic nerve head (ONH). The lower panel shows higher magnification images highlighting RGC staining in each group (Scale bar: 1 mm). B. Quantification of RBPMS+ve cells in control (n = 5), PBS-injected FD (n = 6), and scAAV2-FD-ExSpeU1-eGFP–injected FD (n = 5) mice (superior; *p<0.0003,* inferior; *p<0.0329,* temporal; *p<0.0471* and nasal; *p<0.0172*). Data are presented as mean ± SEM, with each data point representing an individual retina. Statistical analysis was performed using a two-tailed unpaired Student’s t-test with FDR correction. Adjusted p-values are shown: *\*p < 0.05; **p < 0.01*.

### scAAV2-FD-ExSpeU1-eGFP treatment improves visual function in FD mice

To this end, our results demonstrate that intravitreal delivery of scAAV2-FD-ExSpeU1-eGFP significantly improves *ELP1* splicing in the retina, increases RNFL and GCIPL thickness, and prevents RGC loss in FD phenotypic mice. To confirm that rescue of RGC translates to a functional improvement, we conducted a comprehensive assessment of retinal physiology in treated mice. Specifically, we used pattern electroretinography (PERG), a non-invasive, objective method that selectively measures RGC activity^46,47^, to quantify RGC function in *FD* mice injected with scAAV2-FD-ExSpeU1-eGFP. Since we previously observed significant RGC loss by 6 months of age in FD phenotypic mice¹ , we assessed treated and control animals at this time point.

Our findings revealed a marked reduction in PERG amplitude *(Figure 5A)* in FD phenotypic mice compared to controls (p < 0.0062), indicating significant RGC dysfunction. Notably, eyes treated with scAAV2-FD-ExSpeU1-eGFP exhibited significant restoration of PERG amplitude compared to PBS-injected controls (p < 0.0085), demonstrating that ExSpeU1-mediated correction of *ELP1* splicing not only preserves RGC survival but also restores their functional output. These results establish a critical proof-of-concept: molecular rescue of RGCs through splicing correction can lead to functional visual improvement in a mammalian model of FD.

**Figure 5.**
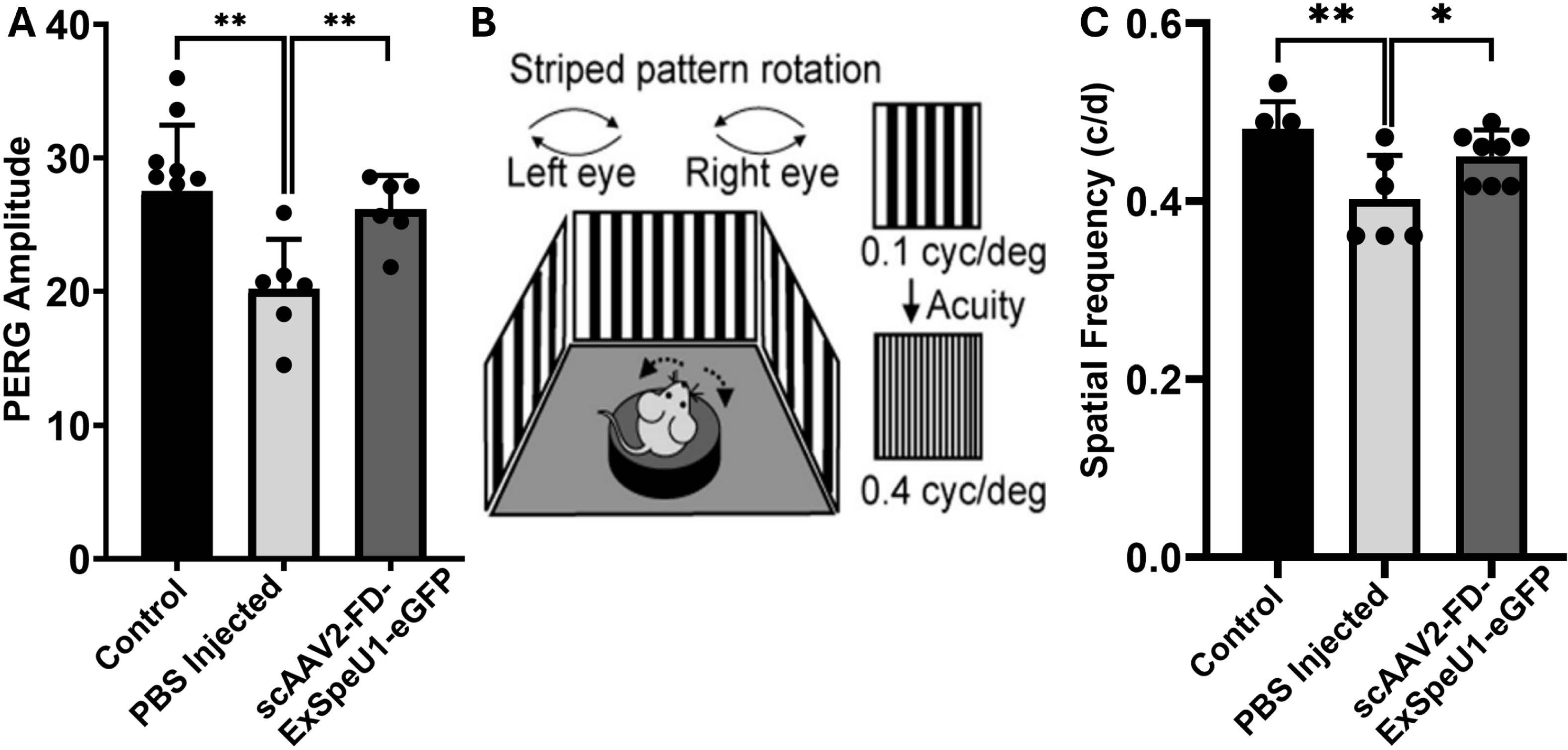
**Rescue of visual function in FD mice**: **A.** Bar graph indicating mean PERG amplitudes of recordings from each FD mouse eye 6 months after intravitreal injection of scAAV2-FD-ExSpeU1-eGFP (*n=6; p<0.0062*) or PBS-injected and control mice (n=8). Significant improvement in PERG amplitude is observed in scAAV2-FD-ExSpeU1*-e*GFP injected eyes compared to PBS-injected eyes (*n=6; p<0.0085*). **B.** A simplified illustration of the OMR assay. High-contrast visual stimulation was applied to track reflexive head movements in response to the rotation of a moving stripe pattern with increased spatial frequency measured as cycles per degree (cyc/deg). **C.** Bar graph representing the OMR of each mouse eye measured as spatial frequency (cycles/degree) in FD mice 6 months after intravitreal injection of scAAV2-FD-ExSpeU1-eGFP (*n=8*; *p<0.0075*) and PBS-injected (n=6) eyes, and control mice (*n=8; p<0.0399*). The data were analyzed with a two-tailed unpaired Student’s *t*-test with FDR correction. Adjusted p-values are shown: *\*p < 0.05; **p < 0.01*

To further evaluate the rescue of visual acuity as an indirect measure of “functional vision” in FD mice injected with scAAV2-FD-ExSpeU1-eGFP and control littermates, we employed an optomotor reflex-based spatial frequency threshold test known as the optomotor response assay (OMR). The OMR assay measures visual acuity by assessing the motor response of mice under photopic conditions to a virtual rotation of stripes of different widths. Mice were able to move freely and were placed on a pedestal located in the center of an area formed by four computer monitors arranged in a quadrangle. The monitors displayed a moving vertical black and white sinusoidal grating pattern *(Figure 5B)*. The results from OMR analysis showed a significant reduction in optomotor reflex in FD mice compared to the control mice (*p<0.0075*). Importantly, a significant improvement in spatial frequency is observed in eyes injected with scAAV2-FD-ExSpeU1-eGFP compared to PBS-injected eyes *(p<0.0399*). *(Figure 5C).* Taken together, these findings demonstrate that FD phenotypic mice exhibit significant visual dysfunction by 6 months of age, and with targeted *ELP1* splicing modulation via FD-ExSpeU1 delivery, effectively restore visual function *in vivo*. Importantly, this is the first demonstration that anatomical rescue of RGCs leads to a measurable improvement in visual function, providing direct evidence that *ELP1* splicing correction can rescue and restore retinal ganglion cell functionality.

## Discussion

FD is a severe neurodegenerative disorder with complex neurological manifestations. Optic neuropathy in FD is a hallmark of the disease, and it resembles well-characterized optic neuropathies such as LHON and DOA^8–10^. Despite progress being made in developing therapies, there is currently no effective treatment for optic neuropathy in FD. All patients suffer from progressive visual decline and will eventually become legally blind by their third decade of life^8–10^. The eye is an easily accessible organ amenable to localized therapeutic intervention, making it ideal for testing the therapeutic efficacy of various treatments using functional, quantifiable parameters. Recent clinical success in mitigating retinal cell loss in retinal dystrophy patients has demonstrated that gene supplementation strategies that introduce a normal copy of a mutant gene can dramatically improve visual function, which led to the first FDA-approved gene therapy drug, Luxturna^48^. These successful endeavors in retinal gene therapy, combined with the recent demonstration that systemic treatment with an SMC rescues the FD phenotype in a mouse model, have provided the foundation for our development of a retina-specific splicing modulatory therapy using ExSpeU1 to provide hope that the drastic visual loss in FD patients can be rescued.

Progressive loss of RGCs and optic nerve degeneration are characteristic features of optic neuropathy in FD. In this study, we showed that intravitreal delivery of FD-ExSpeU1 can correct *ELP1* splicing defects in the retina and rescue the retinal phenotype in FD. Significant rescue of RNFL and GCIPL thickness in the retinas injected with FD-ExSpeU1 at P30 indicates that delivery of FD-ExSpeU1 at postnatal stages is sufficient to prevent RGC death. Our findings are consistent with earlier studies showing that RGC death can be rescued by oral administration of the splicing modulator compound PTC680, even when the treatment started at 3 months of age in a mouse model of FD^49^. This rescue is attributable to the post-natal nature of RGC death in FD. Together, these results support the idea that there is a broad therapeutic window for treating vision loss in FD patients.

Previous studies have shown that RGC loss in FD can be mitigated through splicing modulatory therapies or genetic supplementation with full-length *ELP1* delivered to the retina ^20,21,49^. However, until now, it remained unknown whether preventing RGC loss also translates into functional recovery of these neurons. Splicing modulation or *ELP1* gene supplementation can reduce RGC loss in FD, but it has been unclear whether this preservation restores function. We provide the first direct evidence that *ELP1* splicing modulation not only sustains RGC survival but also restores measurable function, establishing a robust endpoint for therapeutic evaluation. Unlike the well-documented structure–function correlations in glaucoma, such relationships have not been shown in monogenic inherited optic neuropathies like FD^50,51^.

Toxicity induced by AAV-mediated delivery has been one of the significant safety concerns in ocular gene therapy clinical and preclinical trials ^52–54^. This is mainly attributed to the accumulation of overexpressed protein in the target tissues and higher doses of AAV, leading to toxic immunologic responses^55^. Since FD-ExSpeU1 is a small non-coding RNA specifically targeting the mutated splice site, it does not necessarily promote changes in gene expression or alternative splicing^22,25^. Therefore, we expect that ectopic expression of FD-ExSpeU1 will not result in the induction of a significant cellular stress response. In addition, the scAAV2 serotype used in this study predominantly targets the RGC population, which is known to be affected in the FD retina^42,43^. The AAV dose used in our experiments was 1×10^10^ VG total, which is efficient in the correction of splicing defects in the retina. This in vivo dose was informed by in vitro studies where we observed that an MOI of 1,000 vg/cell achieved a near complete *ELP1* splicing correction in FD fibroblasts^45,56^. Given approximately 1 million RGCs in the mouse retina^57^, this dose corresponds to a theoretical MOI of ∼10,000 vg/cell. Adjusting for ∼30% transduction efficiency with intravitreal AAV2 delivery^44^, the effective MOI (∼3,000–3,500 vg/cell) remains within the range shown to be efficacious in vitro, supporting the translational relevance of our dosing strategy. In addition, this dose is comparable to the dosage used in the LHON gene therapy trial (2.4×10^10^VG)^45^, which showed no serious safety concerns in a phase I clinical trial^58^. Even though our data indicate the efficacy of FD-ExSpeU1 in correcting *ELP1* splicing defect and preventing RGC degeneration in FD, we recognize that preclinical dose escalation and safety studies should be performed in the future for the translation of this technology to the clinic.

Previous studies have shown that systemic correction of *ELP1* splicing defects through administration of FD-ExSpeU1 via IP or ICV rescues motor and proprioceptive functions in FD mice^25^. Even though these results are promising, there are several challenges for systemic administration of FD-ExSpeU1 in FD patients. First, systemic administration of AAV in several clinical and preclinical studies has been shown to exhibit hepatotoxicity due to the accumulation of AAV in the liver^59–61^. Accumulation of AAV in dorsal root ganglia (DRG) and associated toxicity is also commonly seen with systemic administration of AAV^62^. Second, the requirement of high vector dosage for systemic administration and associated costs to produce GMP-grade vector pose a major challenge for an ultra-rare disease such as FD^63^. Third, rescue of RGC loss with systemic delivery of FD-ExSpeU1 has not been evaluated. Therefore, there is no clear indication whether systemic delivery of AAV9-FD-ExSpeU1 will be efficacious in rescuing RGC loss in FD. On the other hand, oral delivery of kinetin-derived SMC, such as PTC258 and PTC680, has been shown to rescue RGC loss in FD mouse models ^20,49^. These compounds are currently undergoing toxicology studies to evaluate their safety for potential clinical use. However, no approved treatment currently exists for FD-associated vision loss. With the recent success of retinal gene therapies, there is now a viable and accelerating path to clinical translation for AAV-mediated approaches such as FD-ExSpeU1. In conclusion, our study provides the first evidence that targeted splicing correction using a modified U1 snRNA delivered intravitreally can not only prevent retinal ganglion cell loss but also restore visual function in a mouse model of FD. This functional rescue, achieved through scAAV2-FD-ExSpeU1, is particularly compelling given the precision of the approach and the markedly reduced off-target effects^22,64^. Together, these findings underscore the therapeutic potential of ExSpeU1 for treating FD-associated optic neuropathy and establish a strong rationale for its advancement toward clinical development.

## Data & Materials Availability

All data supporting the findings of this study are included in the published article and its supplementary information files. Additional raw datasets, including imaging files, qPCR data, and statistical analyses, are available from the corresponding authors upon reasonable request. Plasmids encoding scAAV2-FD-ExSpeU1, *ELP1* FD minigene, and related AAV vector backbones can be provided to qualified researchers under a material transfer agreement with Massachusetts General Hospital.

## Ethics Statement

All animal procedures were conducted in compliance with the Association for Research in Vision and Ophthalmology (ARVO) Statement for the Use of Animals in Ophthalmic and Vision Research. Experimental protocols were reviewed and approved by the Institutional Animal Care and Use Committee (IACUC) of Massachusetts General Hospital (protocol number: 2006N000196). All efforts were made to minimize animal suffering and to reduce the number of animals used.

## Author contributions

EM, SA, and AC conceived and designed the study. AC, EM, and LV developed the methodology, and AC, KK, EK, MC, SK, DK, GR, MS and JB performed the investigations. FP, LV, and SA provided resources. AC, EM, and KK curated the data, with analyses and visualizations by AC, KK. SK. AC, KK, SK, and EM drafted the manuscript, and all authors reviewed and edited it. EM, SA, and AC supervised the work, and SA secured the funding.

## Declaration of Interests

EM serves on the scientific advisory board (SAB) of ReviR Therapeutics and is an inventor on an International Patent Application Number PCT/US2021/012103, assigned to Massachusetts General Hospital and PTC Therapeutics, entitled “RNA Splicing Modulation” related to the use of BPN-15477 in modulating splicing.

**Figure S1.** Complete gel image of the FD minigene splicing assay performed in HEK293T cells, demonstrating dose-dependent splicing correction by the scAAV2-FD-ExSpeU1-eGFP vector. Lane M shows the molecular size marker. Lanes 1–3 correspond to cells transfected with 1 µg of vector,,lanes 4–6 with 0.5 µg, lanes 7–9 with 0.25 µg, and lanes 10–12 with 0.125 µg. The final three lanes (13-15) represent cells transfected with the control FD minigene without vector treatment. The aberrantly spliced mutant *ELP1* transcript is detected as the upper band at 550 bp, while the correctly spliced wild-type transcript appears as the lower band at 480 bp. Increasing scAAV2-FD-ExSpeU1-eGFP vector concentration resulted in progressive restoration of the wild-type splicing product.

**Figure S2:** Complete gel picture indicating patient fibroblasts (*ELP1c.2204+6T>C*) infected with different concentrations of scAAV2-FD-ExpeU1-eGFP (Lanes 3,4 and 5 are triplicates of 10^4^ MOI and lanes 6,7 and 8 are triplicates of 10^3^ MOI) showing near complete correction of *ELP1* splicing in FD compared to untransfected control (lanes 1 and 2). Wild-type fibroblasts were included as a control (Lanes 9 and 10)

**Figure S3.** Complete gel image of splicing analysis of human *ELP1* transcripts from the retina of *TgFD9* mice injected with scAAV2-FD-ExSpeU1-eGFP vector (lanes 7-12) or PBS-injected control (lanes 1-6). RT-PCR products reveal the aberrantly spliced mutant transcript at 363 bp and the correctly spliced wild-type transcript at 289 bp. Both PCR products were confirmed by Sanger sequencing.

