## Supplementary figures and images for "AAV2-mediated intravitreal delivery of exon-specific U1 snRNA rescues optic neuropathy in a mouse model of familial dysautonomia"

### Supplementary figure 1

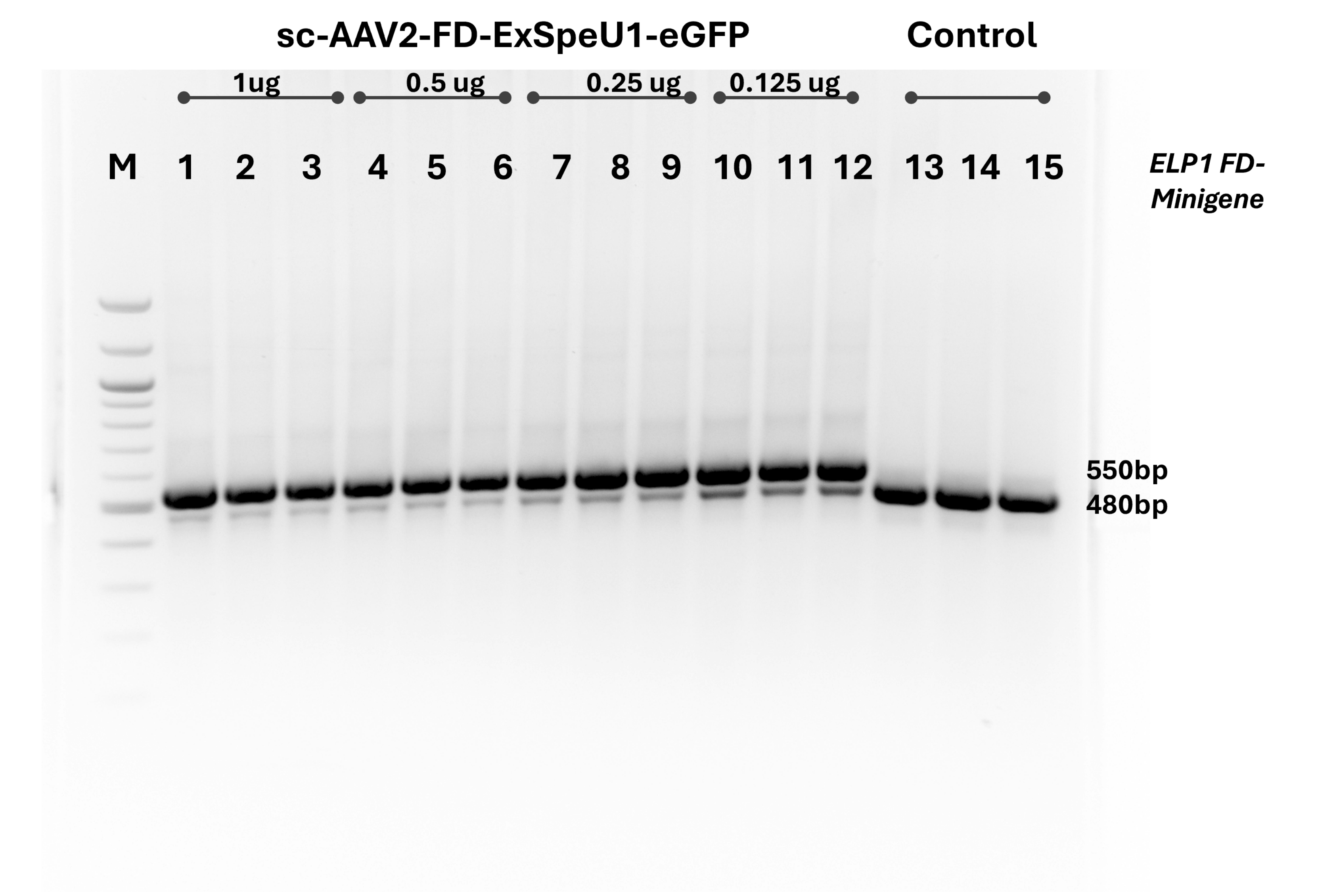

### Supplementary figure 2

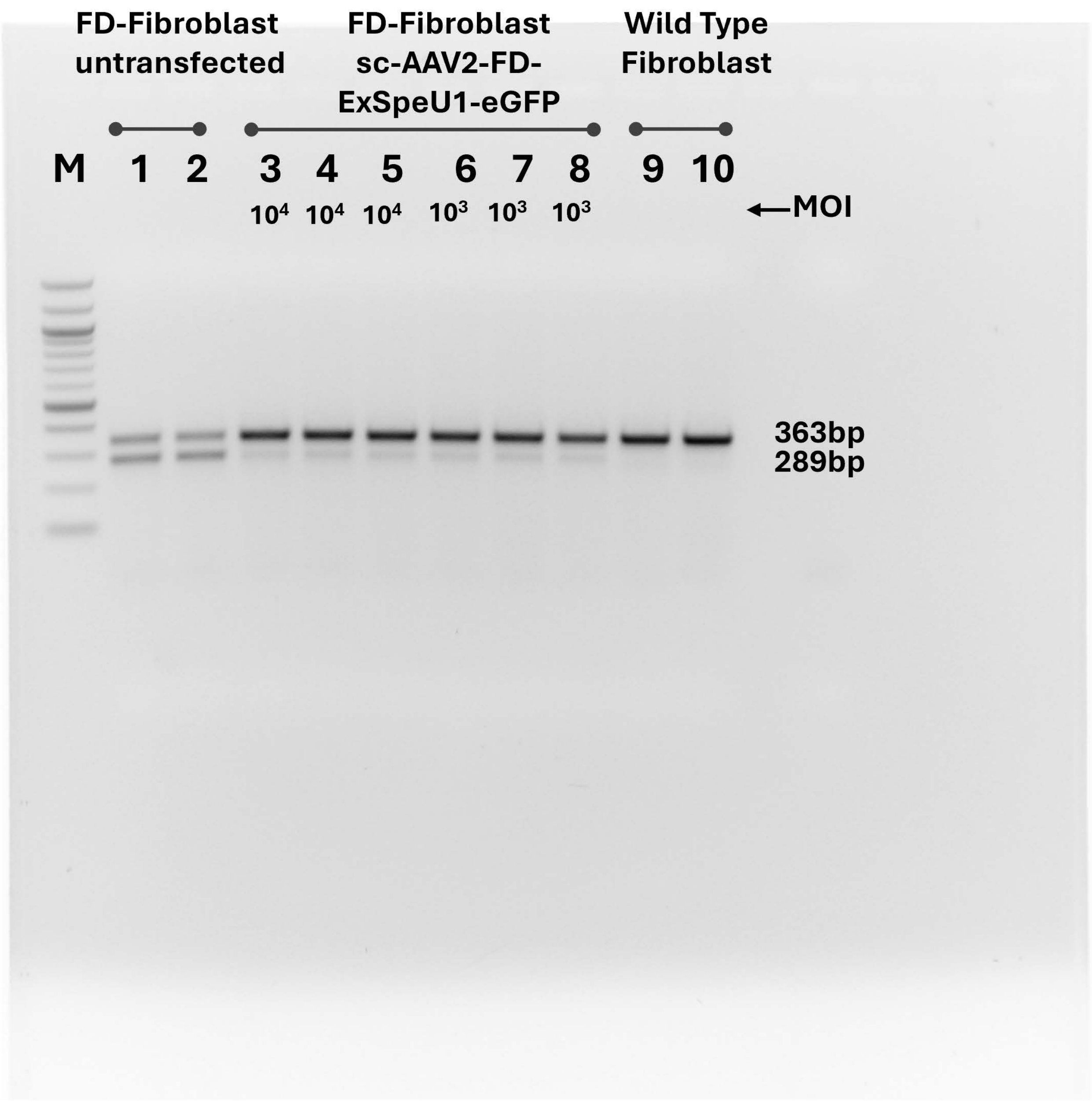

### Supplementary figure 3

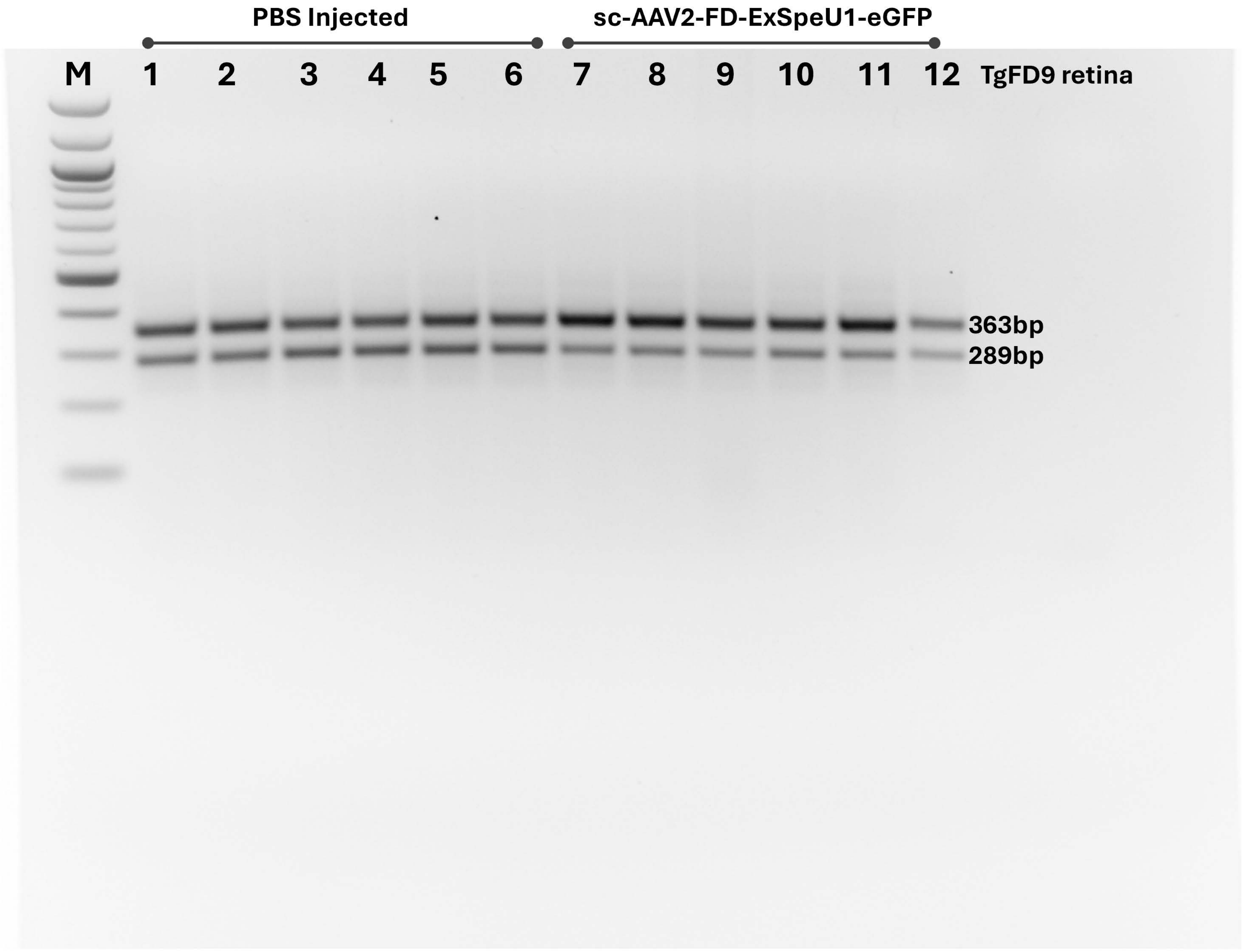
